# Contextual extinction of drug-associated discriminative stimuli fails to attenuate drug-vs-food choice in rhesus monkeys

**DOI:** 10.1101/753699

**Authors:** Matthew L Banks, Blake A. Hutsell, S Stevens Negus

## Abstract

Relapse within the context of a substance use disorder can be triggered by cues that function as discriminative stimuli to signal contingencies of drug availability and promote drug-taking behavior. Extinction procedures can weaken this association between drug-associated cues and drug-taking behavior and may reduce the probability of relapse. This study evaluated a regimen of extinction training on cocaine and heroin self-administration in rhesus monkeys under a drug-vs.-food choice procedure. Behavior was initially maintained under a concurrent schedule of food (1-g food pellets; fixed-ratio 100 schedule) and cocaine injections (0-0.1 mg/kg/injection; fixed-ratio 10) (n=4 males) or heroin injections (0-0.01 mg/kg/injection; fixed-ratio 10) (n=3 females and 1 male) during daily 2-h choice sessions. Subsequently, choice sessions were supplemented by daily 20-h extinction sessions for 14 consecutive days. During extinction sessions, drug-associated discriminative stimuli were presented, but responding produced saline injections. Drug continued to be available during choice sessions. Prior to extinction, both cocaine- and heroin-maintained dose-dependent increases in drug-vs.-food choice. Exposure to 14 extinction sessions failed to significantly decrease drug choice and increase food choice. These preclinical results do not support the effectiveness of extinguishing drug-associated discriminative stimuli as a non-pharmacological treatment strategy for reducing drug choice.

---

One prominent theory of substance use disorder conceptualizes it as a chronic relapsing disorder (Heilig et al., 2021; Leshner, 1997; Volkow et al., 2016). Relapse can be triggered by cues (e.g., the sight of drug paraphernalia) that are known in operant conditioning terms as “discriminative stimuli” and that signal drug availability contingencies to support drug use. Human laboratory (O’Brien et al., 1998; Volkow et al., 2003) and preclinical research (Fredriksson et al., 2021; Madangopal et al., 2019) research has suggested one obstacle to the clinical treatment of substance use disorders is the persistent effectiveness of discriminative stimuli to promote drug-taking behavior even after protracted drug abstinence periods. Modest decreases in cocaine use have been achieved with behavioral therapy implementing extinction training behavioral therapy to blunt the association between cocaine-associated discriminative stimuli and cocaine reinforcement by engaging inhibitory learning processes (Bouton, 1993, 2002; O’Brien et al., 1990). In these extinction-training procedures, subjects are brought to a laboratory environment and exposed to cues that may function as discriminative stimuli in the natural environment, but the subjects are then denied the opportunity to use cocaine. Such extinction training has been shown to reduce both physiological and subjective responses to cocaine-associated discriminative stimuli and produce at least a transient decrease in relapse probability when subjects return to the natural environment (O’Brien et al., 1990). However, a significant clinical challenge is to identify all cues that function as discriminative stimuli in human addicts.

These challenges are more manageable in preclinical research environments, where investigators have more precise control over the identity of discriminative stimuli. Accordingly, the study goal was to determine effects of a 14-day extinction-training regimen on cocaine and heroin self-administration in rhesus monkeys under a drug-vs-food choice procedure that has been extensively used to evaluate other pharmacological and non-pharmacological treatment strategies (Banks, Blough, Fennell, et al., 2013; Banks, Blough, & Negus, 2013; Nader & Woolverton, 1992; Negus, 2005). Rhesus monkeys were initially trained to self-administer cocaine or heroin during daily 2-h “choice” sessions to assess behavioral allocation between intravenous drug injections and an alternative nondrug food reinforcer. Once choice behavior was stable, daily choice sessions continued, and 20-h extinction sessions were introduced from 1200-0800 h each day for 14 consecutive days. During extinction sessions, the drug key was illuminated green for cocaine and yellow for heroin as during choice sessions, but responding produced saline injections. We hypothesized that extinction of responding on the drug-associated key during 20-h extinction sessions would also decrease active drug self-administration during choice sessions and correspondingly increase food-maintained responding. Confirmation of this hypothesis would provide an empirical basis for subsequent studies to optimize extinction-training parameters and to investigate the neurobiological mechanisms that might mediate these extinction effects on drug choice.

## Methods

### Subjects

Four adult male rhesus monkeys (*Macaca mulatta*) of Chinese- or Indonesian-origin with prior cocaine self-administration histories and three adult females and one adult male of Chinese- or Indonesian-origin with prior opioid self-administration histories served as subjects. Monkeys were surgically implanted with a double-lumen catheter (STI Flow, Raleigh, NC) inserted into a major vein. Monkeys weighed 7-13 kg and were maintained on a fresh fruit and food biscuits diet (Lab Diet Monkey Biscuits no. 5045; PMI Nutrition, St. Louis, MO). Water was continuously available via an automatic system. A 12-h light-dark cycle was in effect (lights on from 0600 to 1800 h). Animal research and maintenance were conducted according to the Guide for the Care and Use of Laboratory Animals (National Research Council, 2011). Animal facilities were licensed by the United States Department of Agriculture and accredited by the Association for Assessment and Accreditation of Laboratory Animal Care. The Institutional Animal Care and Use Committee approved the research and enrichment protocols. Monkeys had visual, auditory, and olfactory contact with other monkeys throughout the study. Operant procedures and foraging devices were provided for environmental manipulation and enrichment. Videos were played for additional enrichment.

### Apparatus and catheter maintenance

Monkeys were housed individually in well-ventilated, stainless steel chambers that also served as experimental chambers. Each chamber was equipped with a custom operant panel that contained two horizontally arranged response keys, a pellet dispenser (Model ENV-203-1000; Med Associates, St Albans, VT), and two syringe pumps (Model PHM-108; Med Associates), one for each lumen of the double lumen catheter. One syringe pump (the “self-administration” pump) delivered response-contingent cocaine or heroin injections during choice sessions, and the second syringe pump (the “treatment” pump) delivered both response-contingent saline injections during extinction sessions and non-contingent saline injections at a programmed rate of 0.1 mLs every 20 min. The intravenous (IV) catheter was protected by a tether and jacket system (Lomir Biomedical, Malone, NY). Catheter patency (loss of muscle tone within 10 s) was periodically evaluated by ketamine (3 mg/kg, IV) administration and after a rightward shift in the drug choice dose effect function produced by an experimental manipulation.

### Behavioral Procedure

Initially, monkeys responded during daily 2-h choice sessions (0900 to 1100 h) that consisted of a five-component concurrent schedule of food pellet and IV drug availability. During each component, responses on the left key were reinforced with food (1-g banana-flavored pellet; Test Diets, Richmond, IN) according to a fixed-ratio (FR) 100 schedule, and responses on the right key were reinforced with IV cocaine (0–0.1 mg/kg/injection) or heroin (0-0.01 mg/kg/injection) according to an FR 10 schedule. A response on one key reset the ratio requirement on the alternative key. Each reinforcer delivery was followed by a 3-s timeout during which all stimulus lights were extinguished, and responding had no programmed consequences. During each component in the cocaine choice procedure, the food and cocaine keys were transilluminated with red and green stimulus lights, respectively. During each component in the heroin choice procedure, the food and heroin keys were transilluminated with red and yellow stimulus lights, respectively. Additionally, the stimulus lights for the drug key flashed on and off in 3 s cycles, and longer flashes were associated with larger drug doses. The drug key was continuously illuminated green (cocaine) or yellow (heroin) during availability of the largest drug dose. Furthermore, schedule completion on the food key always produced delivery of a single food pellet, whereas schedule completion on the drug key produced the available unit cocaine dose (0, 0.0032, 0.01, 0.032, and 0.1 mg/kg/injection during components 1–5, respectively) or unit heroin dose (0, 0.00032, 0.001, 0.0032, and 0.01 mg/kg/injection during components 1-5, respectively) by manipulating the injection volume (0, 0.01, 0.03, 0.1, and 0.3ml/injection, respectively). Each component was in effect until 10 total reinforcers were earned or 20 min elapsed, whichever occurred first.

Once drug-vs.-food choice was stable, defined as the smallest unit drug dose that maintained greater than 80% drug choice did not vary by more than 0.5 log units over three consecutive days, experimental conditions were introduced. First, daily extinction-training sessions were introduced to provide conditions during which drug-associated discriminative stimuli that were previously associated with 0.1 mg/kg/injection cocaine or 0.01 mg/kg/injection heroin availability were now associated with IV saline injections for 20 h each day (1200 to 0800 h). During extinction-training sessions, the same right response key associated with drug injections during the choice session was transilluminated green for cocaine or yellow for heroin, and schedule (FR10) completion turned off the stimulus light, activated the treatment pump to deliver saline (0.3 mL/injection), and initiated a 15-min timeout. At the end of the 14-day experimental period, the 20-h extinction sessions were terminated, and choice behavior was monitored for an additional 3 days. Second, saline was substituted for cocaine or heroin both during the 2h drug-vs-food choice session and the 20-h extinction session for 14 consecutive days.

### Data Analysis

The primary dependent measures for choice sessions were percent drug choice, defined as {(number of completed ratios on the drug-associated key ÷ total completed ratios) × 10} and the numbers of total, drug, and food reinforcers earned per session. Data for each of these variables was averaged within a monkey for the last 3 days of each 7-day treatment block, and then between monkeys. Due to equipment failure on extinction-training session day 7 in 3 out of 4 cocaine choice monkeys, results for the 7-day treatment block were expressed as the mean for days 5 and 6. Drug choice results were analyzed using two-way repeated-measures ANOVA with treatment time (7 or 14 days) and drug dose as the fixed effects and significant effects were followed up by a Dunnet post-hoc test. Degrees of freedom were corrected using Geisser-Greenhouse epsilon. T_1/2_ values were calculated using the nonlinear fit function in Prism. Statistical significance criterion was set at the 95% confidence level (P<0.05). Analyses were conducted using Prism 8.0 for Mac (GraphPad Software, La Jolla, CA).

### Drugs

(−)-Cocaine HCl and Heroin HCl were provided by the National Institute on Drug Abuse (Bethesda, MD) Drug Supply Program. Cocaine and heroin were dissolved in sterile saline and passed through a sterile 0.22-μm filter (Millipore Corp, Billerica, MA) before IV delivery. Doses were calculated using the salt form.

## Results

### Extended saline session effects on drug-vs-food choice

Figure 1A and 1B shows extinctions session effects on the cocaine-choice dose-effect function and heroin-choice dose-effect function, respectively. Under baseline conditions, both cocaine and heroin maintained a dose-dependent increase in cocaine choice. Both cocaine and heroin were chosen almost exclusively over food at the largest two drug doses. 14-days of extinction-training sessions failed to significantly attenuate both cocaine and heroin choice as demonstrated by only significant main effects of drug dose, no significant main effect of time or dose × time interactions (cocaine dose: F(1.1,3.2)=60.2, p=0.003; heroin dose: F(1.4,4.1)=42.7, p=0.002). Figure 1C and 1D show extinction-training sessions did not significantly alter total, food, or drug choices during choice sessions. Figures 2 and 3 show individual subject data for extinction session effects on cocaine choice and heroin choice, respectively. 14 days of extinction training resulted in a half-log rightward shift in the cocaine-choice dose-effect function in three out of four monkeys. In contrast, extinction training produced weaker effects on heroin-choice dose-effect functions and in a smaller proportion of individual subjects (i.e., two out of four monkeys). Figure 1E and 1F show rates of saline-maintained responding over the 14 extinction-training sessions. There was a decrease in operant responding over time in both groups (cocaine group: mean T_1/2_ ± standard error = 2.2±0.7 sessions; heroin group: mean T_1/2_ ± standard error = 2.5±0.9 sessions).

**Figure 1:**
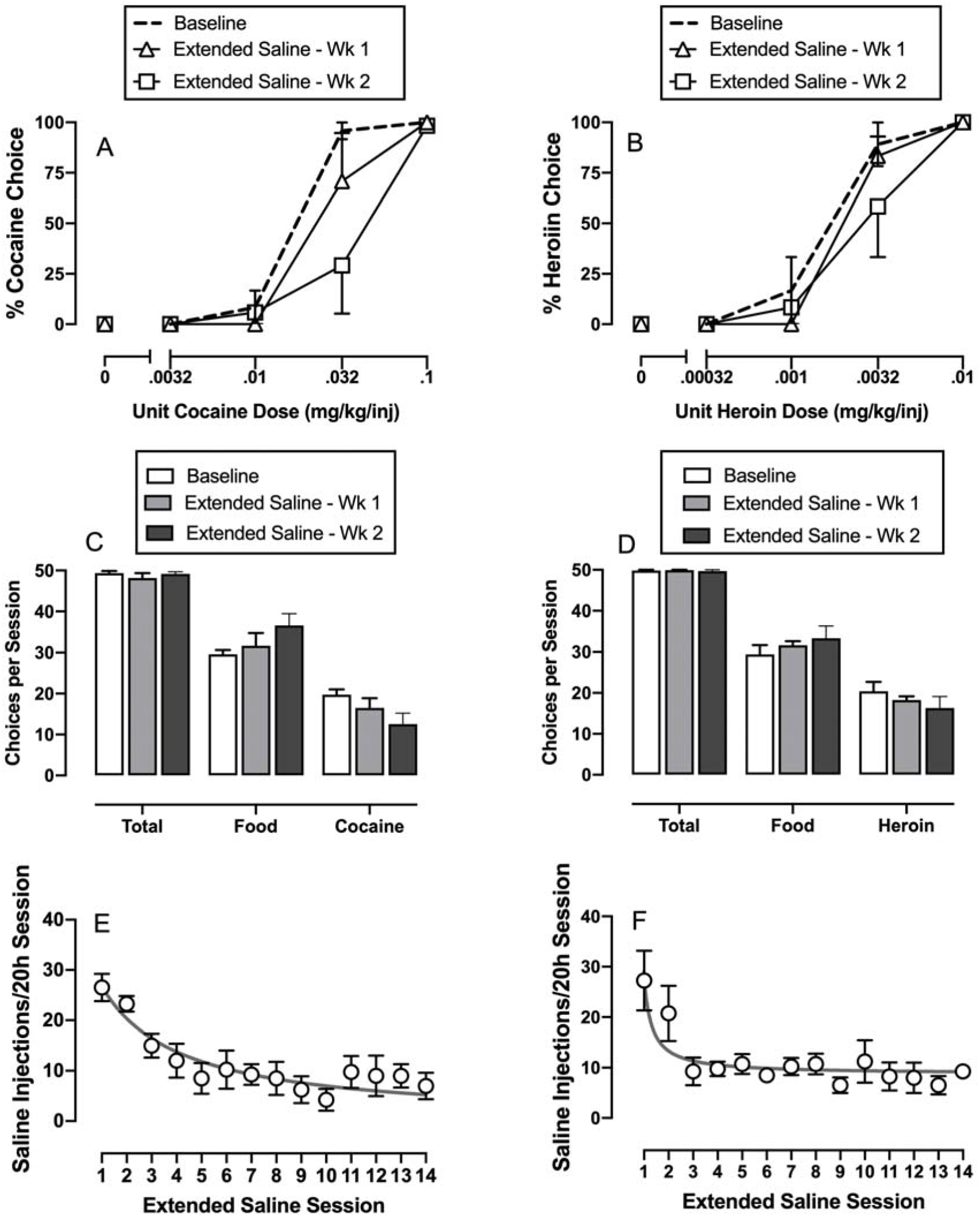
Extended saline session effects on cocaine-vs-food and heroin-vs-food choice. *Note:* Extended saline session effects on (A) cocaine-vs-food choice, (B) heroin-vs-food choice and (C and D) total, food, and drug choices, respectively (A and B) Ordinates: percent drug choice. Abscissa: unit drug dose in milligrams per kilogram per injection. (C and D) Ordinates: total choices summed across each component. Abscissae: total, food, or drug choices. Panels E and F show the time course of the number of saline injections earned during the 20-h extinction training session for cocaine and heroin experiments, respectively. All symbols and bars represent mean ± s.e.m. obtained during days 5-6 (7 day) for cocaine studies or days 5-7 (7 day) for heroin studies and 12-14 (14 day) of the 14-day extinction period in separate groups of four monkeys.

**Figure 2:**
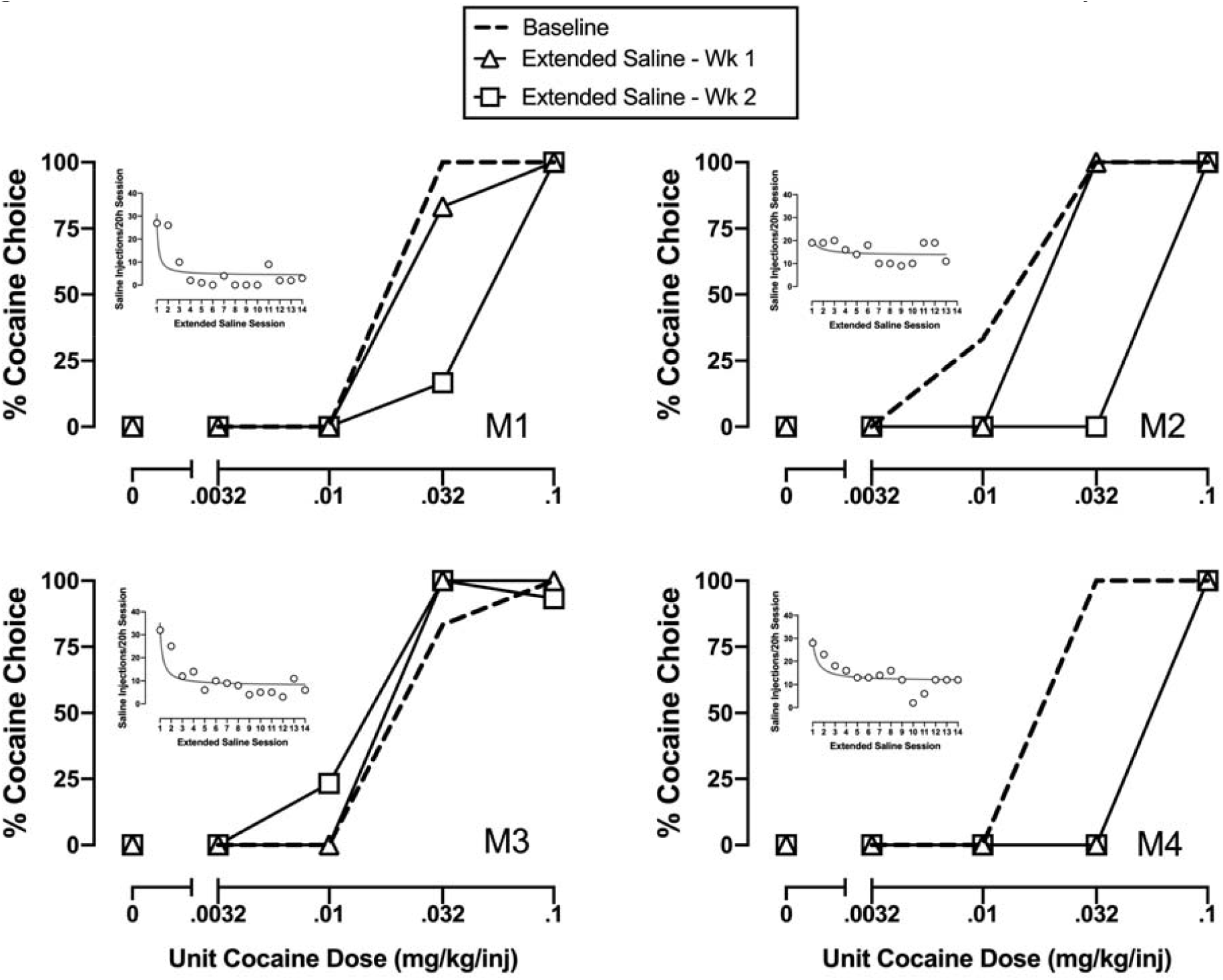
Extended saline session effects on cocaine-vs-food in individual monkeys. *Note:* Extended saline session effects on cocaine-vs-food choice in individual monkeys. Ordinates: percent cocaine choice. Abscissa: unit cocaine dose in milligrams per kilogram per injection. All symbols represent mean ± s.e.m. obtained during days 5-6 (7 day) and 12-14 (14 day) of the 14-day extinction period in each monkey. Insert shows the time course of the number of saline injections earned per 20-h extended saline session over the 14-day extinction period.

**Figure 3:**
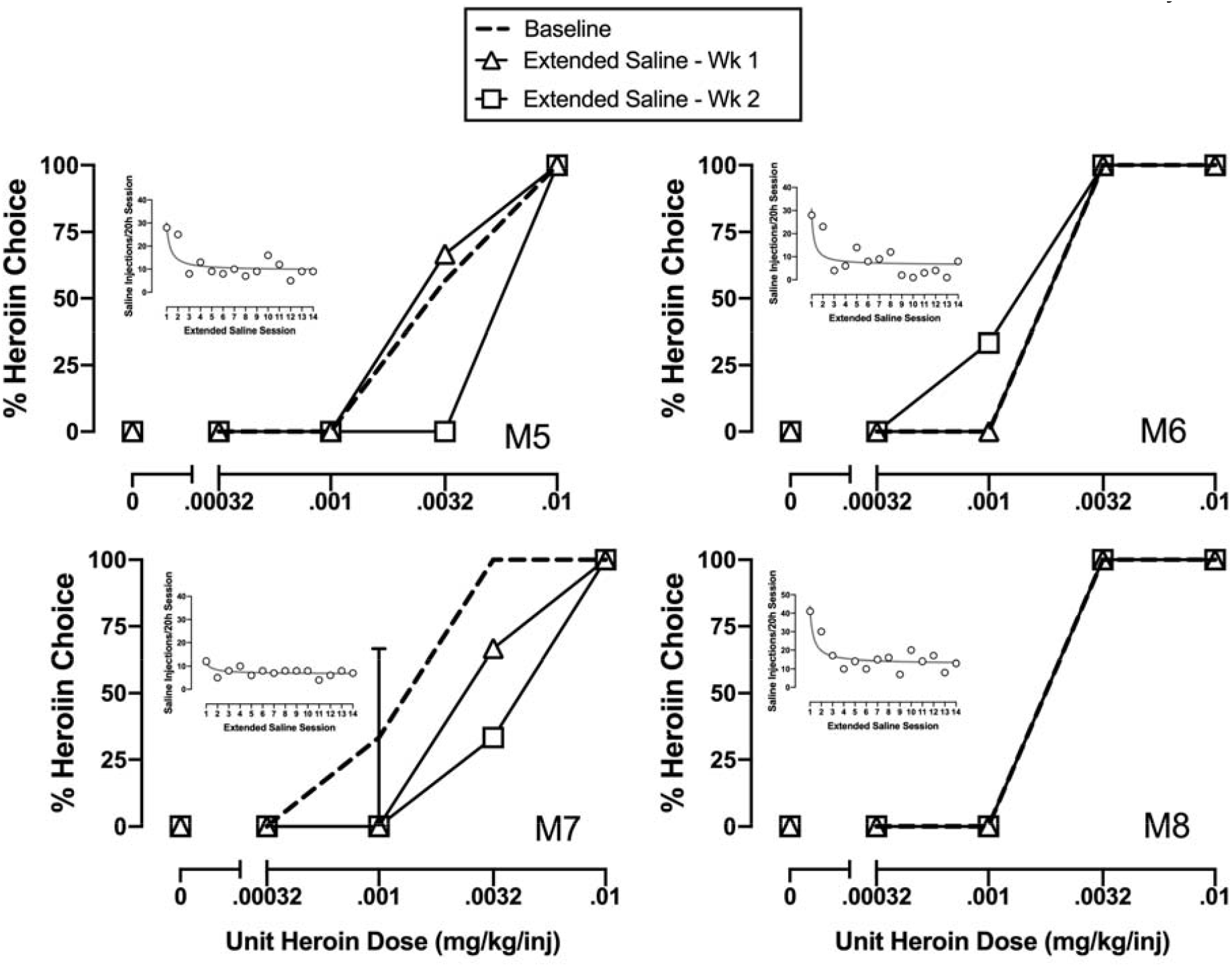
Extended saline session effects on heroin-vs-food choice in individual monkeys. *Note:* Extended saline session effects on heroin-vs-food choice in individual monkeys. Ordinates: percent heroin choice. Abscissa: unit heroin dose in milligrams per kilogram per injection. All symbols represent mean ± s.e.m. obtained during days 5-7 (7 day) and 12-14 (14 day) of the 14-day extinction period in each monkey for the choice data. Insert shows the time course of the number of saline injections earned per 20-h extended saline session over the 14-day extinction period.

### Saline substitution effects on drug-vs-food choice

Figure 4 shows effects of substituting saline during the drug-vs-food choice session in addition to the 20-h extinction-training session. 14 days of saline substitution significantly decreased the percentage of choices completed on the key previously associated with cocaine (choice × time: F(1.9,5.7)=10.9, p=0.012) or heroin (choice × time: F(1.2,2.4)=13.7, p=0.049) compared to components under baseline conditions where IV drug injections were delivered following schedule completion. Figure 4C and 4D show that behavior was reallocated towards food and away from the key previously paired with cocaine or heroin injections during both the first and second weeks of the 14-day saline substitution period (cocaine: dependent measure × time: F(1.3,4)=37.43, p=0.003; heroin: dependent measure × time: F(1,2)=36.11, p=0.027). Figure 4E and 4F show rates of saline-maintained responding over the 14 extinction-training sessions. There was a decrease in operant responding over time (cocaine group: mean T_1/2_ ± standard error = 2.5±0.8 sessions; heroin group: mean T_1/2_ ± standard error = 5.5±3.6 sessions.

**Figure 4:**
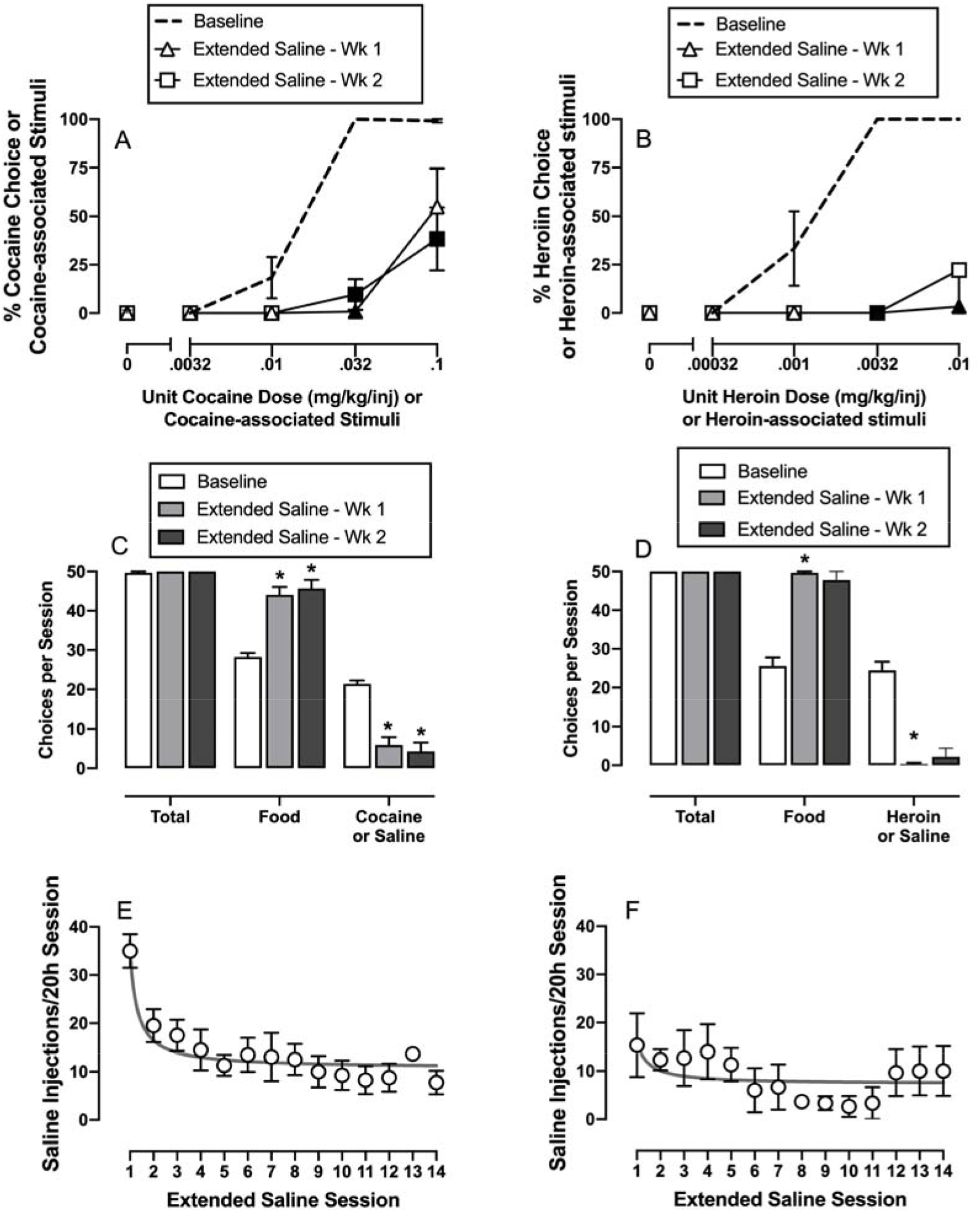
Effects of saline substitution during the drug-vs-food choice session in addition to the 20-h extended saline session on behavior allocation between food and drug-associated stimuli. *Note:* Effects of 14-day saline substitution during the 2h drug-vs-food choice session in addition to the 20h extinction component on behavioral allocation between food and previous drug-associated stimuli in rhesus monkeys. (A and B) Ordinates: percent drug choice or drug-associated stimuli. Abscissa: unit drug dose in milligrams per kilogram per injection or drug-associated stimuli. (C and D) Ordinates: total choices summed across each component. Abscissae: total, food, and drug or saline choices. Panels E and F show the time course of the number of saline injections earned during the 20-h extended saline session. All symbols and bars represent mean ± s.e.m. obtained during days 5-7 (7 day) and 12-14 (14 day) of the 14-day saline substitution period in separate groups of four monkeys for the cocaine studies and three monkeys for the heroin studies. Filled symbols and asterisks denote statistical significance (p<0.05) compared to baseline conditions.

## Discussion

The aim of the present study was to determine the effects of extinguishing behavior occasioned by drug-associated discriminative stimuli outside the drug self-administration session on drug vs. food choice. There were two main findings. First, saline-contingent responding during extinction sessions declined rapidly to stable low levels during 14 extinction session exposure days. Thus, extinction session introduction did produce evidence of behavioral extinction during presentation of drug-associated discriminative stimulus. Second, extinction session exposure also produced a more gradual nonsignificant decrease in cocaine choice and complementary increase in food choice during cocaine choice sessions in three out of four monkeys. Furthermore, this effect was especially prominent during availability of 0.032 mg/kg/injection cocaine demonstrating that an inhibitory learning effect was produced (Bouton, 1993, 2002). Extinction session exposure was both less effective and occurred in a smaller number of monkeys responding for heroin choice. Taken together, these results provide weak preclinical evidence that extinction of responding occasioned by a drug-associated discriminative stimulus in one context can decrease drug self-administration in a different context.

Both cocaine- and heroin-maintained dose-dependent increases in choice over a nondrug food alternative reinforcer in rhesus monkeys. The present results are consistent with the extant literature using both nonhuman primates (Czoty et al., 2005; Foltin et al., 2015; Nader & Woolverton, 1991; Woolverton & Balster, 1979) and rats (Beckmann et al., 2019; Thomsen et al., 2013; Townsend, Schwienteck, et al., 2021) demonstrating the feasibility of establishing drug choice over a nondrug alternative reinforcer. Moreover, these preclinical data are consistent with human laboratory studies demonstrating dose-dependent drug-vs-money choice for both cocaine and opioids (Brandt et al., 2020; Hart et al., 2000; Lile et al., 2016). Furthermore, the reproducibility of this baseline drug-choice behavior in monkeys provides an empirical framework for interpreting both nonpharmacological and pharmacological manipulations on drug choice.

Extinction training is a process of learning new stimulus-response contingencies such that discriminative stimuli previously paired with consequent stimuli delivery (e.g., IV drug infusion) no longer predict the availability of that consequent stimulus. The present results during the 20h extinction sessions and the saline substitution during both the choice and 20h extinction sessions are consistent with this hypothesis and provide empirical evidence that extinction learning can be modeled in the preclinical research setting consistent with the extant preclinical literature (for reviews, see Chesworth & Corbit, 2017; Conklin & Tiffany, 2002). The present results also extend this preclinical literature to the evaluation of extinction training on drug self-administration in a drug-vs-food choice context. To the degree to which extinction training had any effect on drug-vs-food choice, reasons for the qualitative different effects of extinction training on cocaine choice vs. heroin choice are not presently clear.

The effectiveness of extinction training on drug choice can also be compared to other nonpharmacological and pharmacological manipulations. For example, nonpharmacological manipulations such as increasing drug price, increasing the magnitude of the alternative nondrug reinforcer, and punishing cocaine choice all result in decreases of cocaine choice that were of similar magnitude as the present extinction training results (Banks, Blough, & Negus, 2013; Chow & Beckmann, 2020; Johnson et al., 2016; Nader & Woolverton, 1991; Negus, 2003; Negus, 2005; Thomsen et al., 2013). Furthermore, the magnitude of the extinction training effect on cocaine choice in the present study is similar to effects of amphetamine maintenance on cocaine choice in monkeys (Banks, Blough, Negus, 2013a; Banks, Blough, et al., 2013b; Moerke et al., 2017) and rats (Thomsen et al., 2013). However, two caveats when interpreting the present results are worth noting. First, the extinction training effect on cocaine choice was slower to develop compared to both nonpharmacological and pharmacological treatment effects described above. Second, the extinction effect on both cocaine and heroin choice was smaller in magnitude than Food and Drug Administration-approved pharmacological treatments such as naltrexone and buprenorphine maintenance on opioid choice in monkeys (Townsend, Bremer, et al., 2021; Townsend et al., 2020) and rats (Townsend et al., 2019). Thus, the small extinction effect on cocaine choice in the present study should be interpreted within the broader context of clinically approved treatment strategies for substance use disorders and highlight the need for further research to determine whether extinction training represents a clinically-viable nonpharmacological treatment approach either alone or in combination with pharmacological approaches.

The present cocaine choice results are generally consistent with results from the limited clinical literature evaluating extinction training. For example, a small outpatient study that reported the extinction treatment groups had more weeks with clean urines than the nonextinction groups (O’Brien et al., 1990). In nontreament-seeking smokers, denicotinized cigarettes in the absence of nicotine replacement therapy decreased rates of craving, smoking rates, and the motivation to smoke during treatment (Donny et al., 2007). However, nicotine replacement therapy has been demonstrated to be more effective in promoting abstinence than extinction training (Rose, 2006). Depot naltrexone maintenance effects on opioid-taking behavior has been described in terms of extinction learning as the potential behavioral mechanism (Nunes et al., 2020). A key difference between naltrexone maintenance and the extinction experiments in the present study and those by O’Brien (1990) is the environmental context under which extinction training occurs. Although the housing chamber was the same for both the drug-vs-food choice sessions and extinction sessions in the present study, there were contextual differences in the environment where extinction of behavior occurred such as the absence of the food discriminative stimulus for food pellets during the extinction sessions and the time of day that might have weakened the effectiveness of the extinction sessions. In contrast, naltrexone maintenance results in extinction of behavior in the natural or laboratory environment where opioid use occurred. This later strategy appears to be more effective in facilitating new learning of operant contingencies that result in decreased drug-taking behavior and increased behavior maintained by nondrug alternatives.

